# *De novo* genome assembly of *Myricaria laxiflora* provides insights into its molecular mechanisms under flooding stress

**DOI:** 10.1101/2024.03.09.584251

**Authors:** Xiang Weibo, Li Linbao, Huang Guiyun, Dun Bicheng, Chen Huiyuan, Ma Xiaobo, Zhang Haibo, Xiao Zhiqiang, Liu Jihong, Yang Zhen, Wu Di

**Author notes:** Weibo Xiang and Linbao Li contributed equally to this work.

## Abstract

*M. laxiflora* is an endangered plant distributed in the Yangtze River floodplain of China, which often suffers from flood hazards during its growth and development. Due to the lack of a reference genome for *M. laxiflora*, the molecular regulatory mechanism of its waterlogging stress remains unclear. Here, we report for the first time the high-quality reference genome of *M. laxiflora* by using HiFi sequencing and Hi-C. The total assembly size of the *M. laxiflora* genome was 1.29 Gb with a scaffold N50 size of 29.5 Mb. A total of 23,666 genes encoding proteins and 5,457 ncRNA were predicted in this reference genome. In the *M. laxiflora* genome, a total of 101 MylAP2/ERF genes were identified and divided into 5 subgroups. Many MeJA responsive elements, ABA responsive elements, and hypoxia responsive elements were found in the promoter regions of these MylAP2/ERF genes. Based on transcriptome data analysis, a total of 74 MylAP2/ERF genes can respond to flooding stress. Meanwhile, it was found that three genes (MylAP2/ERF49/78/91) which belong to the same branch with RAP2.2 gene exhibited different expression trends under flooding stress. Our results provide valuable information on the molecular regulatory mechanism of flooding stress in *M. laxiflora*.

## 1. Introduction

*Myricaria laxiflora* (Tamaricaceae, 2n=24) is a perennial deciduous shrub belonging to the family Tamaricaceae [1,2]. It is a typical rare plant in the riparian zone, mainly distributed in the middle and upper reaches of the Yangtze River in China [1]. *M. laxiflora* is a flood tolerant plant that sleeps in summer and autumn, reproduces in winter and spring, and grows on gravel beaches in the Yangtze River water level rise and fall zone [3,4]. During the flood season of the Yangtze River every year, the *M. laxiflora* hibernate underwater, and when it reaches the dry season, they begin to rapidly grow and reproduce [2]. The construction of the Three Gorges Dam (TGD) has seriously threatened the native habitat environment of *M. laxiflora* [5]. During the summer when *M. laxiflora* are affected by floods, they will prevent plant death by regulating the total soluble sugar, fructose, and starch content in the body. *M. laxiflora* can be an ideal plant for studying the molecular mechanism of plant flooding tolerance [6,7]. In recent years, many studies have been conducted to investigate the effects of flooding stress on the physiological response of *M. laxiflora* [2,8,9]. However, little is known about the molecular mechanism of waterlogging tolerance in *M. laxiflora*, partly due to a lack of high-quality reference genome.

In recent years, significant progress has been made in the molecular regulatory mechanisms of plant waterlogging tolerance [10,11,12]. The response of plants to waterlogging stress is sequentially completed through processes such as signal transduction substance induction synthesis, metabolic adaptation, and morphological adaptation (formation of aerated tissue, adventitious roots, etc.), including induction of low oxygen proteins, transcription factors, ethylene synthesis, production and clearance of reactive oxygen species (ROS), etc [13,14,15,16]. Some plant hormones have been proven to be involved in plant adaptation to flood stress, including ethylene, gibberellin, and auxin [17]. The early signal (ethylene) production can be detected within a few hours after plants are subjected to waterlogging stress, indicating that ethylene plays an important role in regulating plant waterlogging tolerance [10, 17]. Under waterlogging stress, the expression level of ACS (ACC synthase), as the coding gene for ethylene synthesis precursor ACC (1-aminocyclopropane-1-carboxylate), is significantly induced [18,19,20,21]. In Arabidopsis, waterlogging stress induces the expression of AtACS5, AtACS7, and AtACS8 [20]. The expression of OsACS5 gene in rice is also induced by waterlogging stress [19]. After being subjected to waterlogging stress, the ethylene precursor ACC produced in plant roots is transported to the stem through the xylem and then oxidized to ethylene, thereby mediating the plant’s waterlogging tolerance [21].

The transcription factors known to play important roles in response to waterlogging stress include MYB, bZIP, bHLH, and ERF [22]. Among these transcription factors, the regulation of plant waterlogging tolerance mechanism by ERF transcription factors has been extensively studied. The rice ERF transcription factors SK1 and SK2 are induced to express under waterlogging stress, activating the gibberellin (GA) pathway to promote rice internode elongation and growth, thereby enabling normal growth and development of rice under waterlogging stress [23]. Overexpression of the ERF transcription factor *Sub1A* in waterlogging sensitive rice varieties can significantly enhance the waterlogging tolerance of transgenic lines [24,25]. The expression of TaERFVI.1 in wheat is also induced by waterlogging stress [26,27]. Overexpression of TaERFVI.1 can significantly improve the waterlogging tolerance of wheat, but has no adverse effect on yield [27]. The ZmEREB180 transcription factor in maize enhances its waterlogging tolerance by promoting the growth of adventitious roots [28]. Overexpression of the ERF transcription factor *RAP2.2* in Arabidopsis can enable it to adapt to flooding stress [29]. And, the other ERF members such as HRE1 and HRE2 have the similar function as RAP2.2 in Arabidopsis [30]. These results demonstrate that ERF transcription factors play an important role in regulating plant flood tolerance response [31,32,33]. Due to the lack of reference genome information, the role of ERF transcription factors in flood tolerance is still unclear in *M. laxiflora*.

In this study, a high-quality chromosome level reference genome was assembled for *M.laxiflora* at the first time. This reference genome has a genome size of 1.29 Gb, a heterozygosity rate of 0.65%, and a GC content of 35%. a total of 23,666 protein coding genes were predicted, and 23,631 genes were found on the assembled 12 chromosomes, accounting for as high as 99.8%. In addition, a total of 101 MylAP2/ERF transcription factors were identified within the reference genome. And, there are 74 MylAP2/ERF genes showed significant changes in expression abundance under flooding stress in *M.laxiflora*. The results of this study provide value information for the research of *M.laxiflora* in the future.

## 2. Results

### 2.1 The *M.laxiflora* Genome-survey, sequencing, and assembly

The variety (LS7) of *M.laxiflora* comes from the Yangtze River in China and exhibits good tolerance to flood stress. To avoid sample contamination, one of the branches was selected for cutting to obtain the asexual seedlings of LS7 (Figure 1A). The root tips, leaves, stems, flowers, fruits, and roots of LS7 seedlings were collected for genome assembly. To obtain the high-quality genome, we first evaluated its chromosome number and genome size. The results show that *M.laxiflora* is diploid with 24 chromosomes (Figure 1B). According to K-mer (K=17) analysis, the estimated genome size is 1.26Gb, with a heterozygosity of 0.65% and a repeat sequence proportion of 69.89%. Afterwards, a total of 49.2Gb (∼44.7×) of HiFi data was generated, the amount of data is sufficient to meet the assembly of the genome. Meanwhile, GC depth analysis was performed on the HiFi data, which showed a concentrated distribution of GC content, indicating that there was no contamination in the sequencing samples. The preliminary assembly of contig using HiFi data showed that the size of contig was 1.32Gb, the length of N50 was 29.5Mb, and the continuity of contig was good. Finally, a total of 155Gb of Hi-C data was obtained, with an effective Hi-C reads of 77.4Gb (∼64×). The contigs were scaffolded at the chromosome level by using the Hi-C data. A total of 618 scaffolds in the 12 chromosomes were generated and the assembly size of *M.laxiflora* genome was 1.29Gb. As shown in Hi-C heatmap, there is enrichment of intra chromosomal interactions without significant assembly errors (Figure 1C). To evaluate the genome integrity, the Benchmarking Universal Single-Copy Orthologs (BUSCO) analysis was conducted and the results showed that the integrity of this genome was 93%, which suggested the assemblyed *M.laxiflora* genome had a good integrity. Meanwhile, the uniform coverage of the 12 chromosomes indicates good genome assembly. These results indicate that we have obtained a high-quality reference genome for *M.laxiflora*.

**Figure 1.**
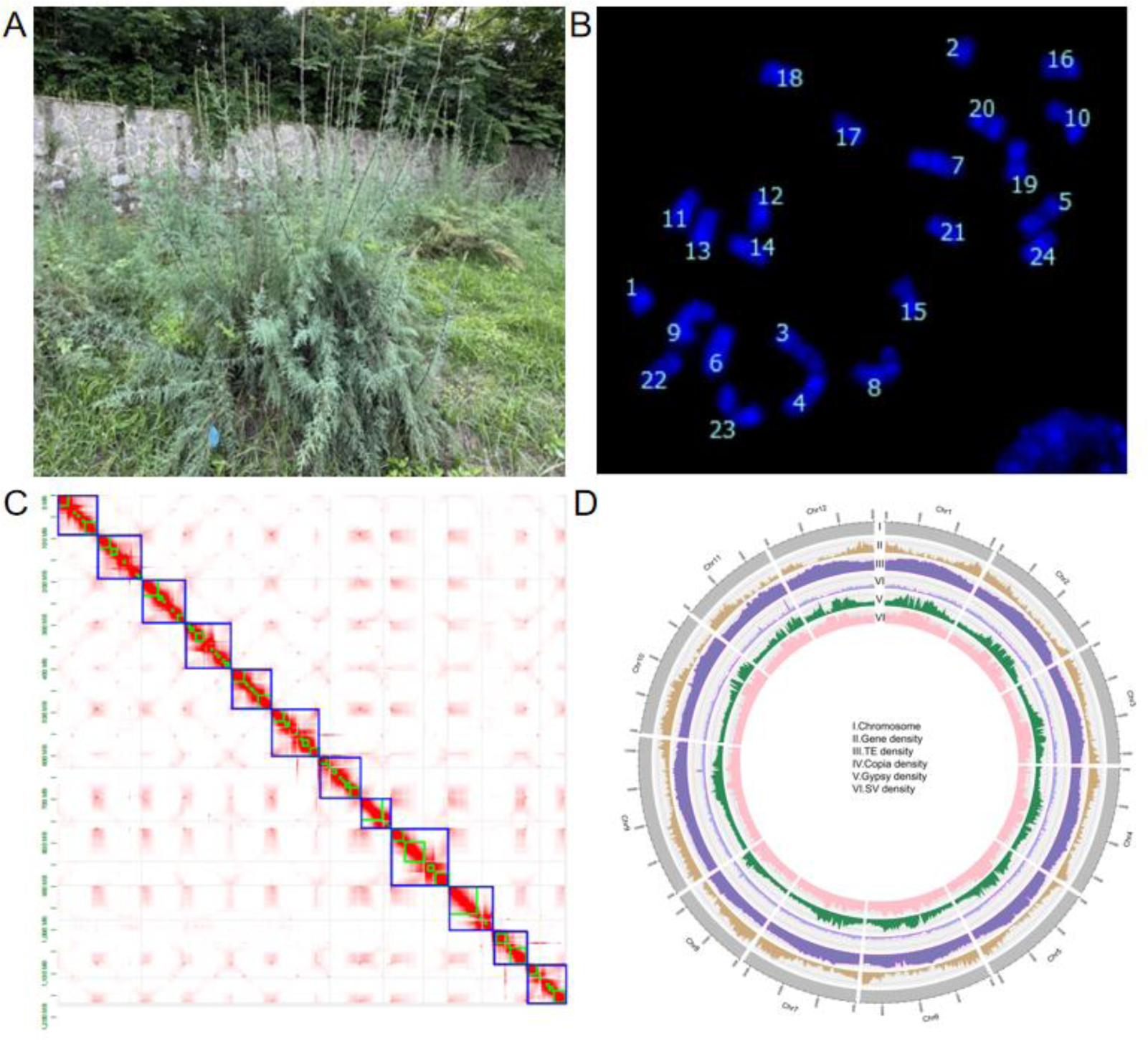
A. Genome sequencing plant(LS7) of *M.laxiflora.* B. Karyotype analysis plots. C. Hi-C heatmap. D. Circular representation of the pseudo-chromosomes.

### 2.2 Genome annotation

This study used multiple methods to annotation the genome of *M.laxiflora*, including ab initio prediction, homology based prediction, and RNA seq based prediction. The results showed that a total of 23,666 protein coding genes were predicted, and 23,631 genes were found on the assembled 12 chromosomes, accounting for as high as 99.8%. The average length of these genes is 6,996.15 bp, and the average exon length is 378.23 bp. The predicted genes of 20,691 genes (87.43%) were annotated from five public databases, including NR, KOG, GO, KEGG, and TrEMBL. Among all annotated genes, 1,270 genes were identified as transcription factors (TFs), accounting for 5.37% of all genes, including bHLH (124), MYB (111), and AP2/ERF (101) families. In addition, we also conducted similarity search and annotation of non-coding RNA (ncRNA) genes, generating a total of 5,457 ncRNA genes, including 94 microRNA (miRNA) genes, 566 transfer RNA (tRNA) genes, 2,648 ribosomal RNA (rRNA) genes, 953 small nucleolar RNA (snoRNA) genes, and 214 small nuclear RNA (snRNA) genes.

A total of 996.66 Mb repetitive sequences (77.16% of the total sequences) were identified through comprehensive methods, including homology based and de novo prediction. Among these repetitive sequences, LTR (long terminal repeat) is the most abundant repetitive element, accounting for 28.04% of the total genome sequence. Most LTRs are Gypsy (21.76%) and Copia (4.50%) components. The amplification of Gypsy transposons is a key driving factor in the genome evolution of *M.laxiflora*. In addition, we found that the complete LTR insertion event in the genome of *M.laxiflora* is a continuous process, with most insertions occurring over the past 4 million years (Figure 2). In addition, a total of 397,683 simple sequence repeats were identified, which will provide useful molecular markers for the molecular breeding of *M.laxiflora*.

**Figure 2.**
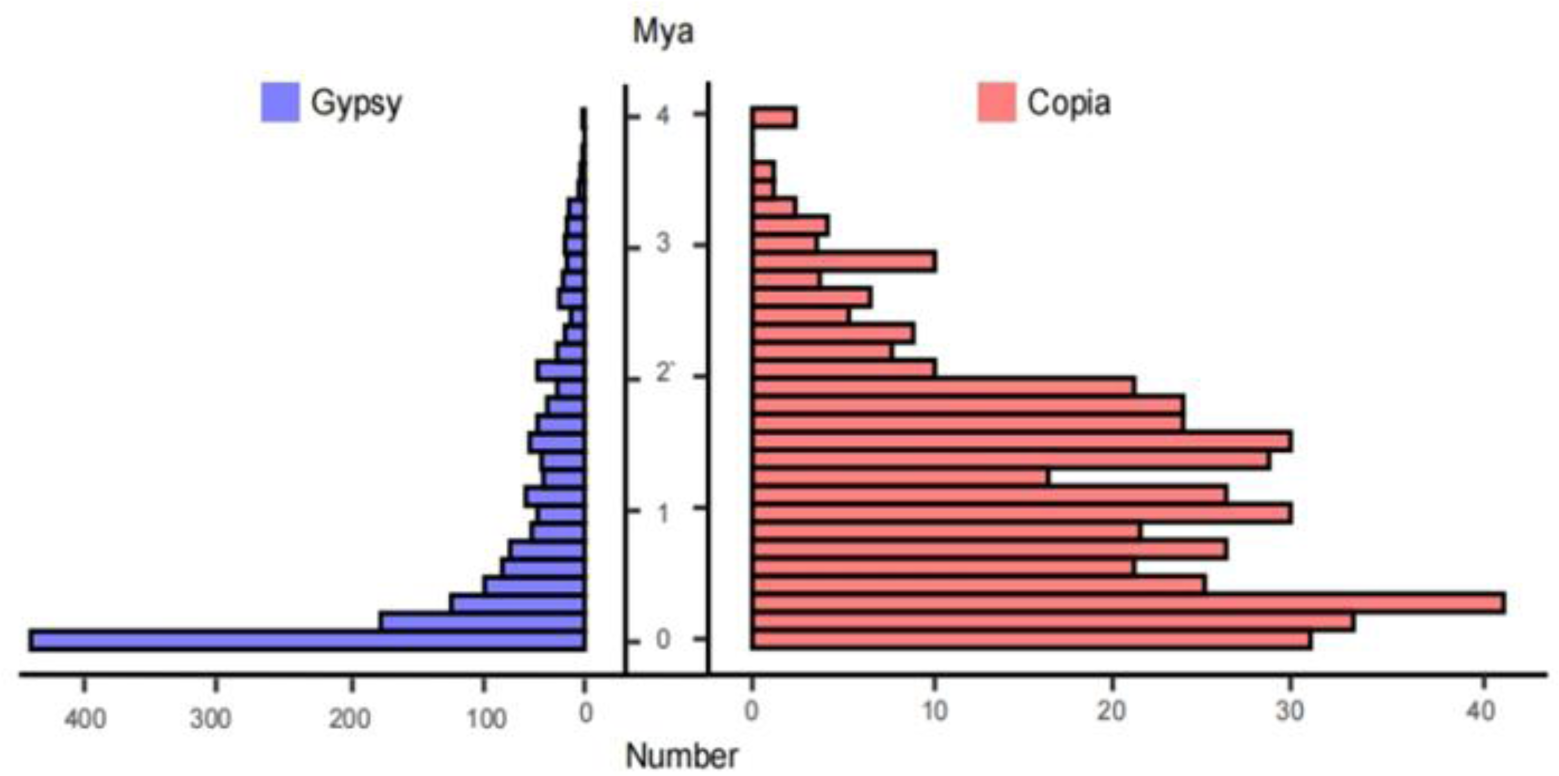
Insertion time distributions of Gypsy and Copia LTR elements.

### 2.3 AP2/ERF gene family identified in the *M.laxiflora* genome

In order to deepen the research on the waterlogging tolerance of *M.laxiflora*, we identified and screened the AP2/ERF gene family in this reference genome through HMM and domain analysis. A total of 101 AP2/ERF genes were found in *M.laxiflora* genome. These 101 members are unevenly distributed on 12 chromosomes of the *M.laxiflora* genome (Figure 3). Among them, the MylAP2/ERF genes are most distributed on chr 9, with a maximum of 21 members, while the MylAP2/ERF genes are least distributed on chr 11, with only 3 members. Afterwards, we renamed the AP2/ERF genes as MylAP2/ERF1-MylAP2/ERF101 based on their positional order on the chromosome. Besides this, we also found that only 100 MylAP2/ERF genes were located on the chromosome, and one gene (MylAP2/ERF101) was not located on the chromosome.

**Figure 3.**
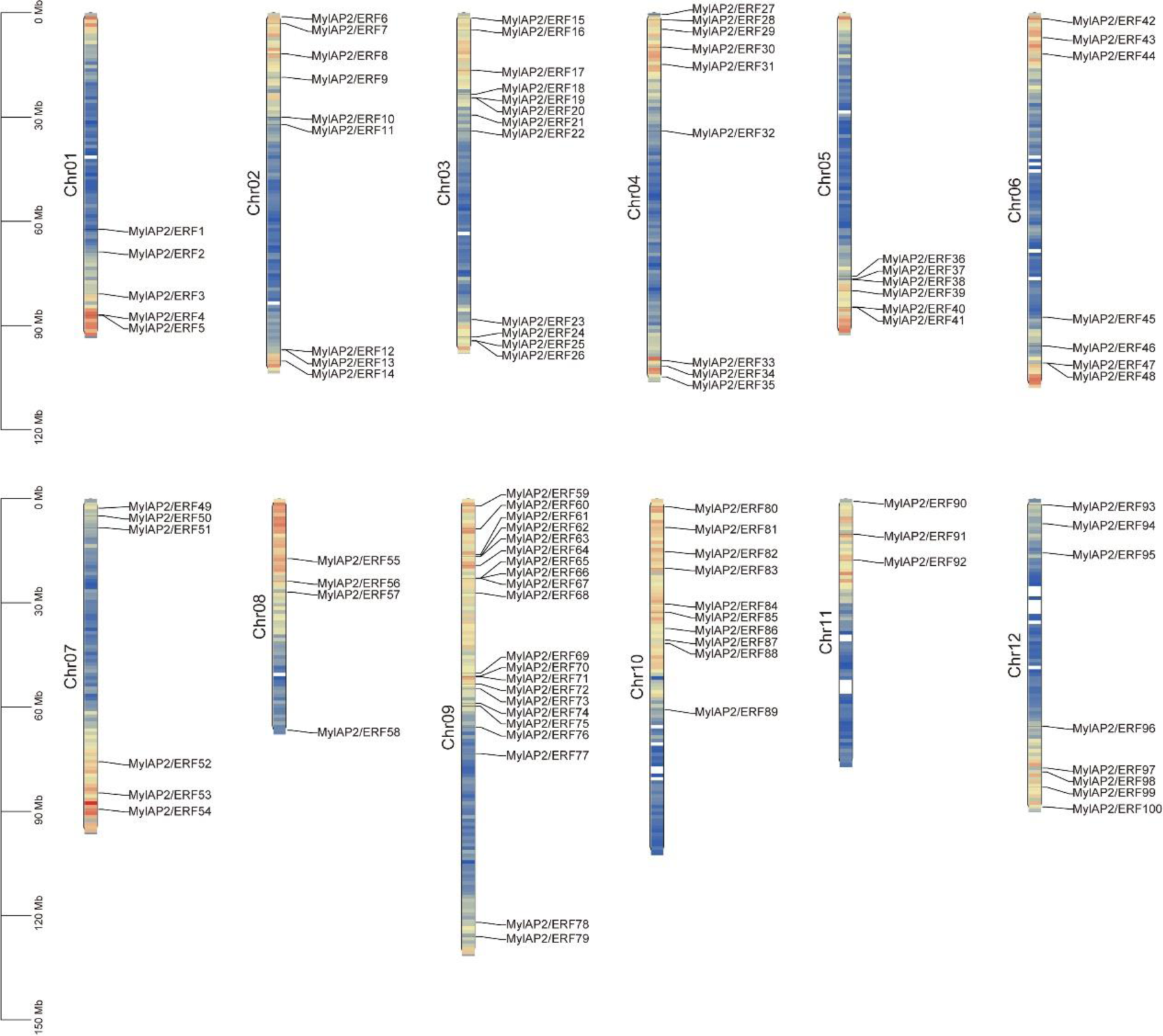
Distribution of AP2/ERF genes on 12 chromosomes of the *M.laxiflora* genome.

### 2.4 Phylogenetic analysis and motif analysis of the MylAP2/ERFs

In order to elucidate the evolutionary relationship of the AP2/ERF gene family between *M.laxiflora* and Arabidopsis which is the model plant, a phylogenetic tree was constructed by using MEGA 11 software. The phylogenetic tree includes 101 MylAP2/ERFs and 142 AtAP2/ERF members. 243 AP2/ERF proteins were assigned to five subgroups, namely AP2, Soloist, RAV, ERF, and DREB subgroups (Figure 4). The ERF subgroup has the highest number of members among the five subgroups, with 104 members, including 64 AtAP2/ERFs and 40 MylAP2/ERFs. In the Soloist subgroup, there are the fewest members, with only 2 AP2/ERF members, including 1 AtAP2/ERF and 1 MylAP2/ERF. The AP2 subgroup consists of 17 AtAP2/ERFs and 17 MylAP2/ERFs, totaling 34 AP2/ERF members. The DREB subgroup consists of 94 AP2/ERFs members, including 54 AtAP2/ERFs and 40 MylAP2/ERFs. The RAV subgroup consists of 6 AtAP2/ERFs and 3 MylAP2/ERFs, with a total of 9 AP2/ERF members. *RAP2.2* (AT3G14230), *RAP2.12* (AT1G53910), *RAP2.3* (AT3G16770), *HRE1* (AT1G72360), and *HRE2* (AT2G47520) have been shown to play important roles in plant flooding stress in Arabidopsis. Interestingly, we found that MylAP2/ERF49, MylAP2/ERF78, and MylAP2/ERF91 genes clustered together with these genes, indicating that they have similar functions in flooding stress in *M.laxiflora*.

**Figure 4.**
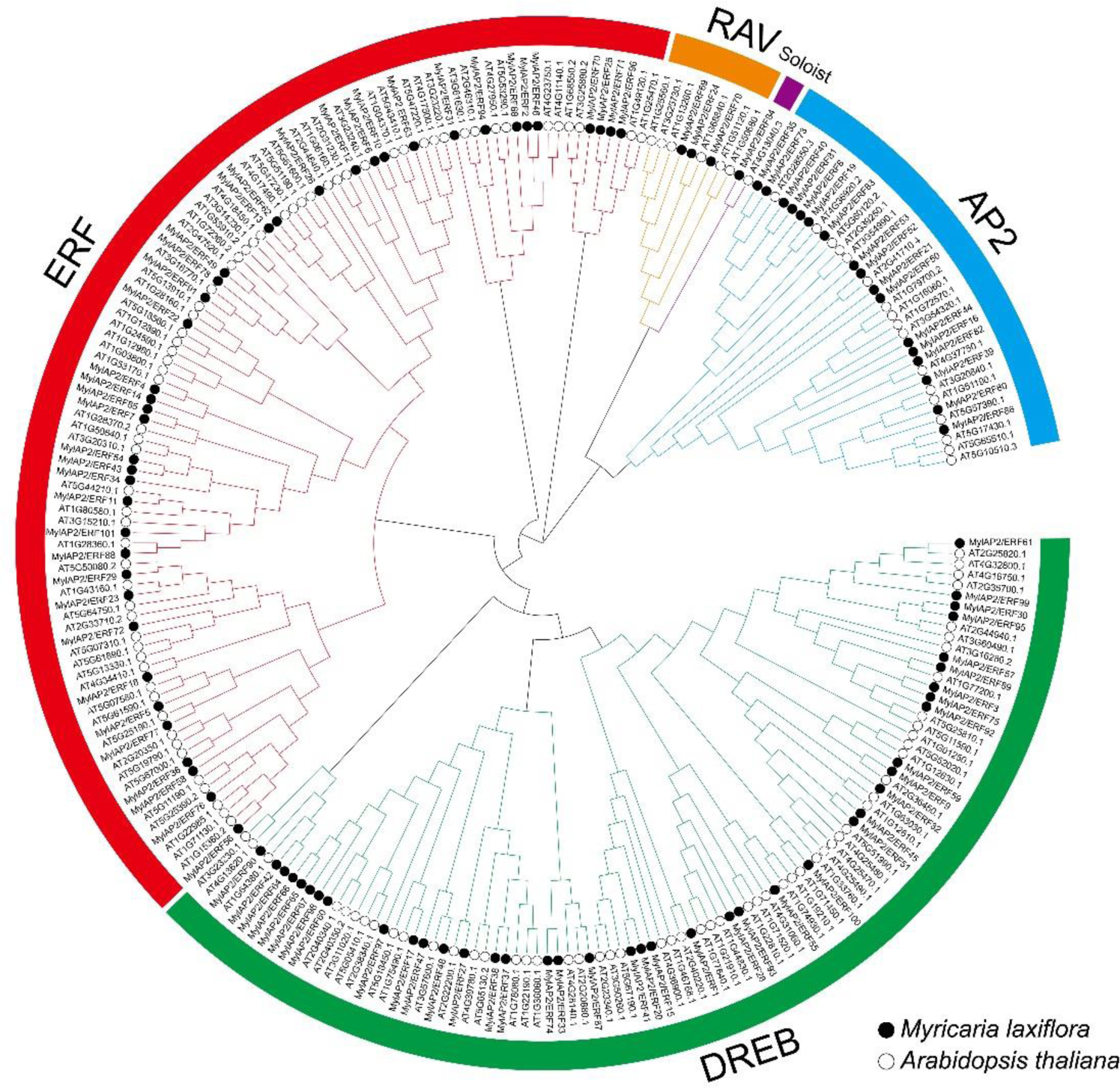
The phylogenetic tree of the AP2/ERF gene family from *M.laxiflora* and *Arabidopsis*.

To identify the conserved motifs of MylAP2/ERFs, the MEME online tool was used for analysis. A total of 10 different conserved motifs were found in the 101 MylAP2/ERF members. The result showed that the sequence length of the 10 motifs ranges from a minimum of 11 amino acids (Motif 2, Motif 3, and Motif 10) to a maximum of 50 amino acids (Motif 4). Some motifs are conserved among MylAP2/ERFs members, while others are unique to only a few MylAP2/ERFs members (Figure 5). For example, Motif1, Motif2, and Motif3 are present in most MylAP2/ERFs, with 98, 99, and 95 MylAP2/ERF genes, indicating that these three motifs are the most important components of the MylAP2/ERFs protein. However, Motif4, Motif6, Motif7, Motif8, and Motif10 are mainly present in the AP2 subgroup members, Motif5 is mainly present in the ERF subgroup, and Motif9 is only present in the DREB subgroup. These results suggest that these motifs may be conserved sequences in different subgroups of MylAP2/ERFs. In addition, the gene structures of 101 MylAP2/ERFs members were analyzed, and the results showed that the gene structures which in the same subgroup were very similar (Figure 5).

**Figure 5.**
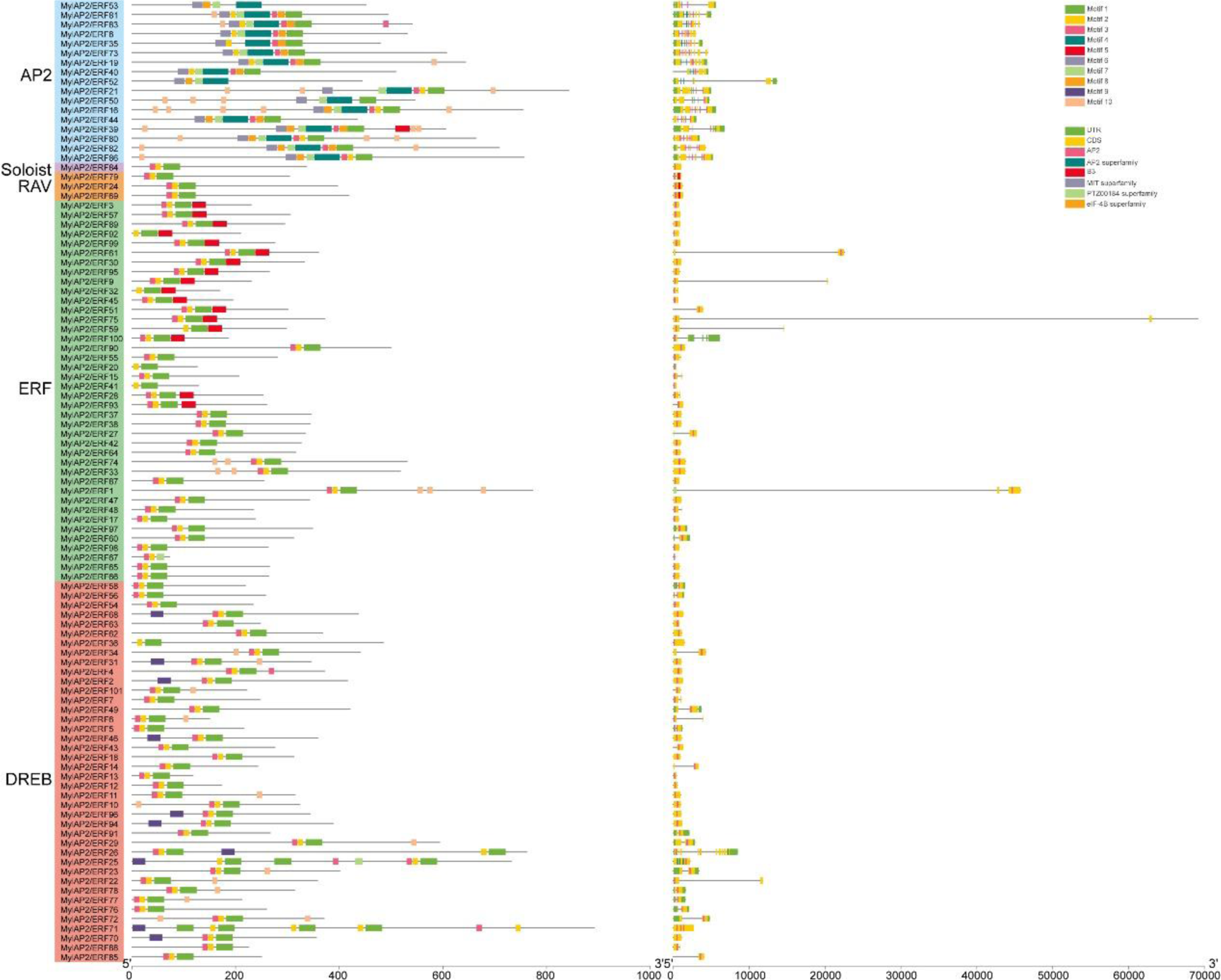
The conserved motifs of MylAP2/ERFs from *M.laxiflora*.

### 2.5 Gene replication analysis among the MylAP2/ERF genes

To investigate possible gene replication events between MylAP2/ERF genes, we localized the predicted MylAP2/ERF using assembled genome data. The results showed that a total of 10 tandem repeat events were found between MylAP2/ERF members, involving 21 MylAP2/ERF genes (Figure 6). For example, there is a tandem repeat event between MylAP2/ERF4 and MylAP2/ERF5 located on chr 1. In addition, 32 fragment duplication events were found, involving 47 MylAP2/ERF genes, such as fragment duplication events between MylAP2/ERF2 located on chromosome 1 and MylAP2/ERF46 located on chromosome 6 (Figure 6). These results indicate that repetitive events are widely involved in the evolution of MylAP2/ERFs.

**Figure 6.**
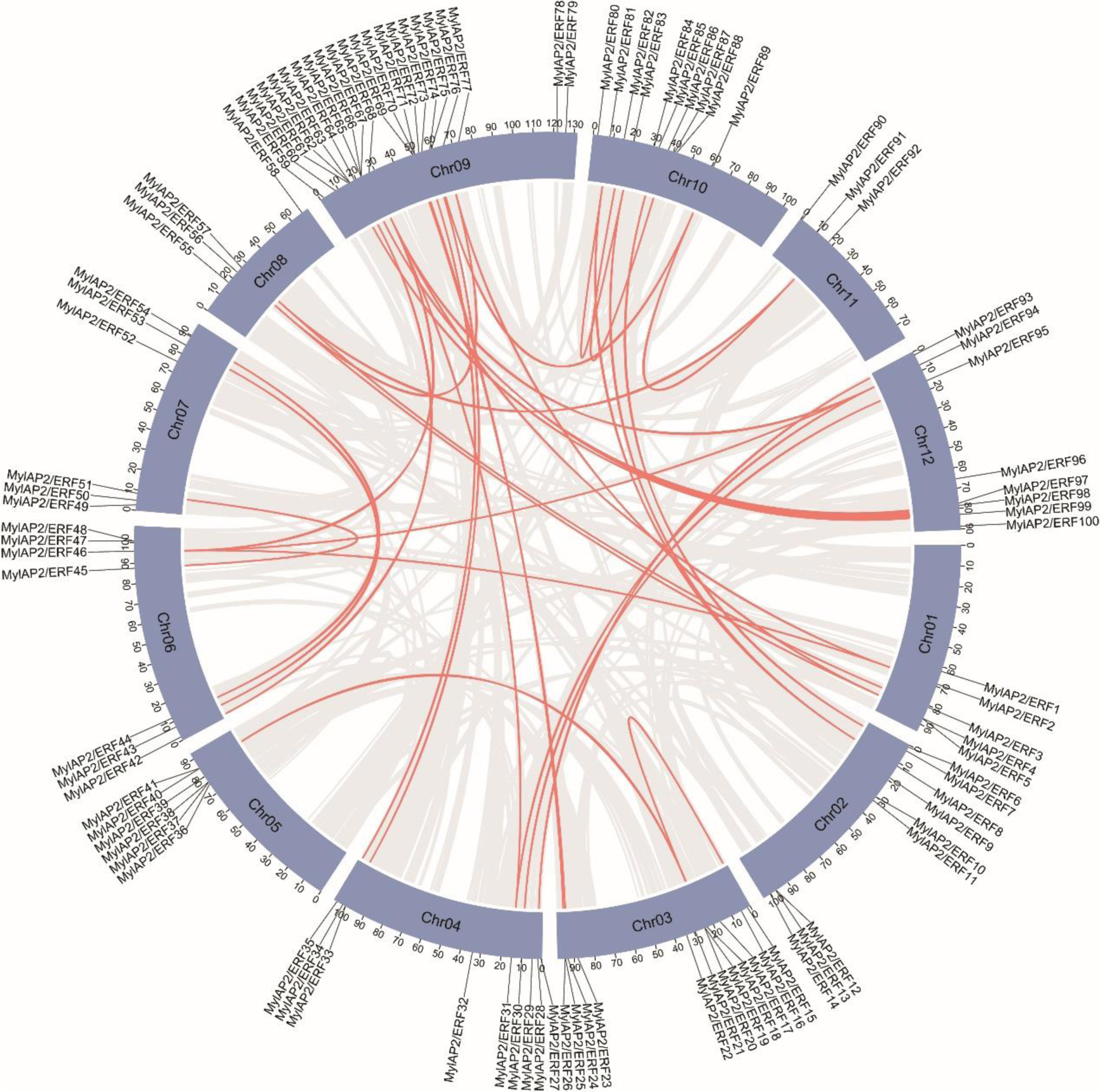
Gene replication events between MylAP2/ERF genes from *M.laxiflora*.

### 2.6 Analysis of *cis*-acting elements in the promoter region of MylAP2/ERFs

In order to further explore the functional potential of the MylAP2/ERFs gene, the PlantCARE database was used to analyze the *cis*-acting elements in the upstream 2000 bp promoter region of the start codon (ATG) of the 101 MylAP2/ERF genes. The results showed that a total of 15 important *cis*-acting elements were discovered, including zein metabolism regulation, abscisic acid response, salicylic acid response, low-temperature response, anaerobic induction, drought response, auxin response, gibberellin response, methyl jasmonate response, defense, and stress response, seed specific regulation, hypoxia specific induction, root specific regulation, flavonoid biosynthesis, and trauma response (Figure 7). Among the 101 MylAP2/ERF genes, the maximum number of *cis*-acting elements predicted on MylAP2/ERF45 was 28, while only one *cis*-acting element was predicted on MylAP2/ERF59. In addition, it was found that the top three *cis*-acting elements among the 101 MylAP2/ERF genes were 248 methyl jasmonate (MeJA) responsive elements, 224 abscisic acid (ABA) responsive elements, and 188 anaerobic induced responsive elements. These results indicate that these MylAP2/ERF genes may be widely involved in the anaerobic stress response process.

**Figure 7.**
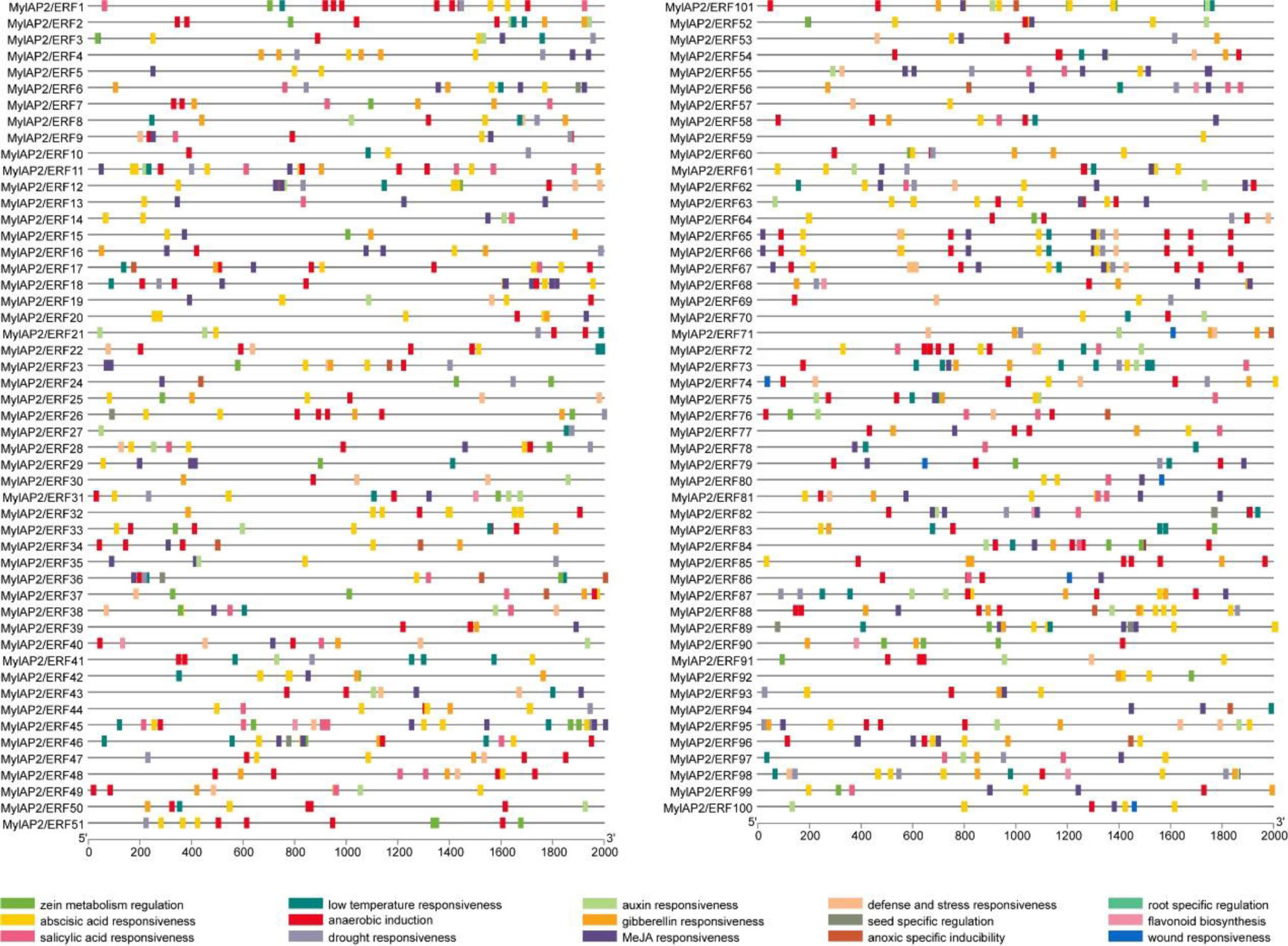
*Cis*-acting elements in the promoter region of MylAP2/ERFs from *M.laxiflora*.

### 2.7 Expression patterns of MylAP2/ERF members in different tissues

In order to clarify the expression patterns of MylAP2/ERF gene family members in different tissues, we obtained expression level data from four tissues of *M.laxiflora*, including stem, leaf, flower, and fruit. As shown in the figure 8, out of 101 MylAP2/ERFs members, 96 genes were expressed in the above 4 tissues, while 5 genes were not expressed. Unexpressed genes may be expressed in other specific tissues or specifically under certain biotic or abiotic stresses. Among the 96 expressed genes, 2, 2, 8, and 84 members are expressed in 1, 2, 3, and 4 different tissues, respectively. For example, MylAP2/ERF6 is only expressed in fruits, while MylAP2/ERF79 is only expressed in leaves, MylAP2/ERF82 is expressed in stems and fruits, and MylAP2/ERF87 is expressed in fruits and flowers. These results indicate that MylAP2/ERF can participate in the growth and development of water cypress branches at different stages through its expression in different tissues.

**Figure 8.**
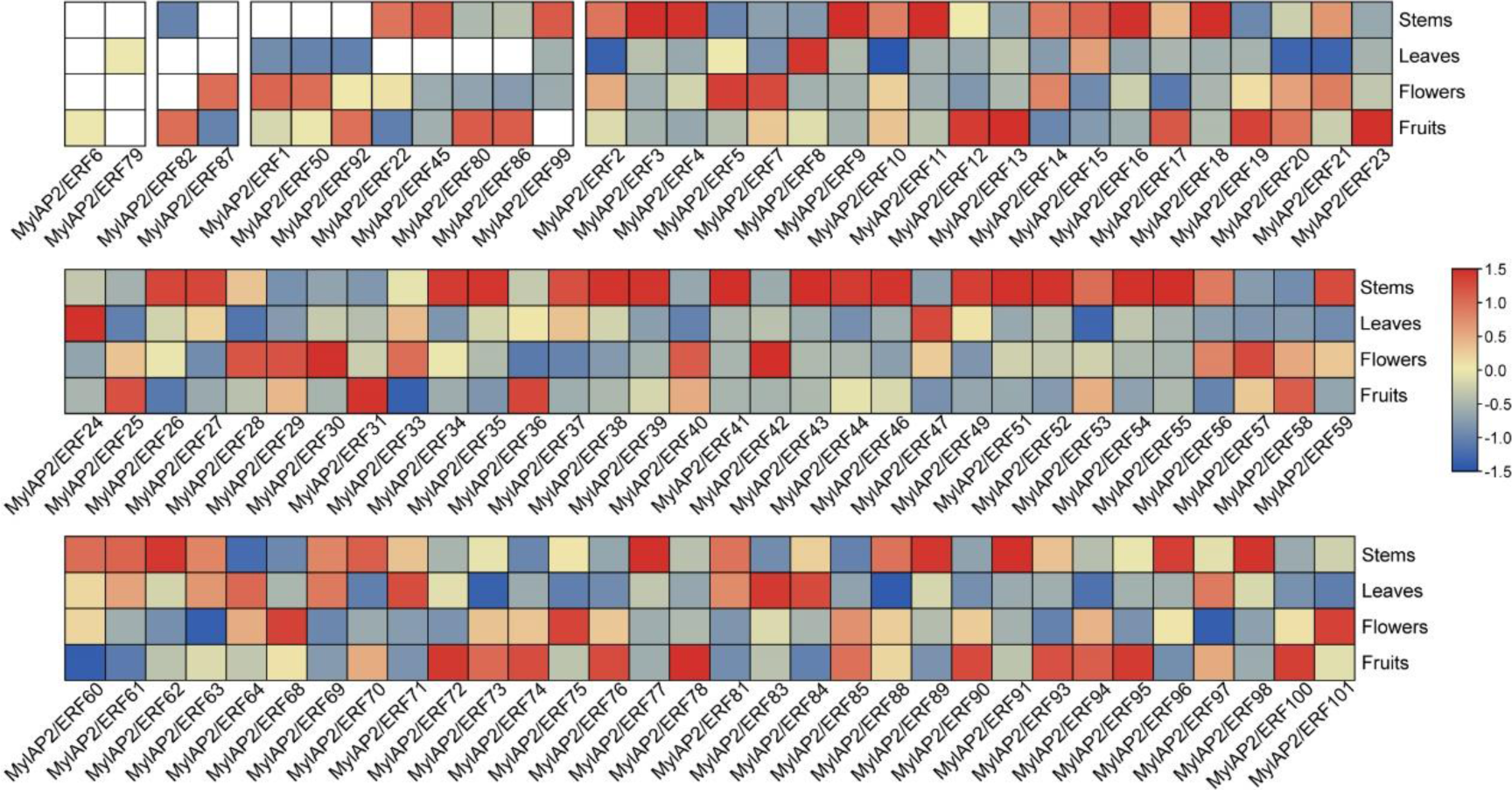
Expression patterns of MylAP2/ERF members in different tissues.

### 2.8 Analysis of expression patterns of MylAP2/ERF in water flooding stress

In order to further clarify whether members of the MylAP2/ERF family responded to flooding stress, transcriptome data of *M.laxiflora* at different time points under flooding treatment were downloaded from the public database. As shown in the figure 9, there are 74 MylAP2/ERF genes showed significant changes in expression abundance under flooding stress at different points. Based on the expression trends at different time points under flooding stress, these 74 genes are classified into 15 different profiles. Among the 15 different profiles, there are three major profiles (50 MylAP2/ERF genes, P <0.05). The expression abundance of the 23 MylAP2/ERF genes which belong to the Profile 0 continues to decrease with the prolongation of waterlogging stress treatment time. The abundance of gene expression which located in Profile 19 (9 MylAP2/ERF genes) continues to increase with the prolongation of waterlogging stress treatment time. The abundance of gene expression which belong to the Profile 16 (8 MylAP2/ERF genes) shows a trend of first increasing, then remaining unchanged, and finally decreasing. The expression abundance of MylAP2/ERF49 and MylAP2/ERF78 showed a trend of first increasing and then decreasing with the increase of flooding stress time (Profile 17), while the expression abundance of MylAP2/ERF91 (Profile 18) showed a pattern of first increasing, then decreasing, and finally increasing again with the increase of flooding stress time.

**Figure 9.**
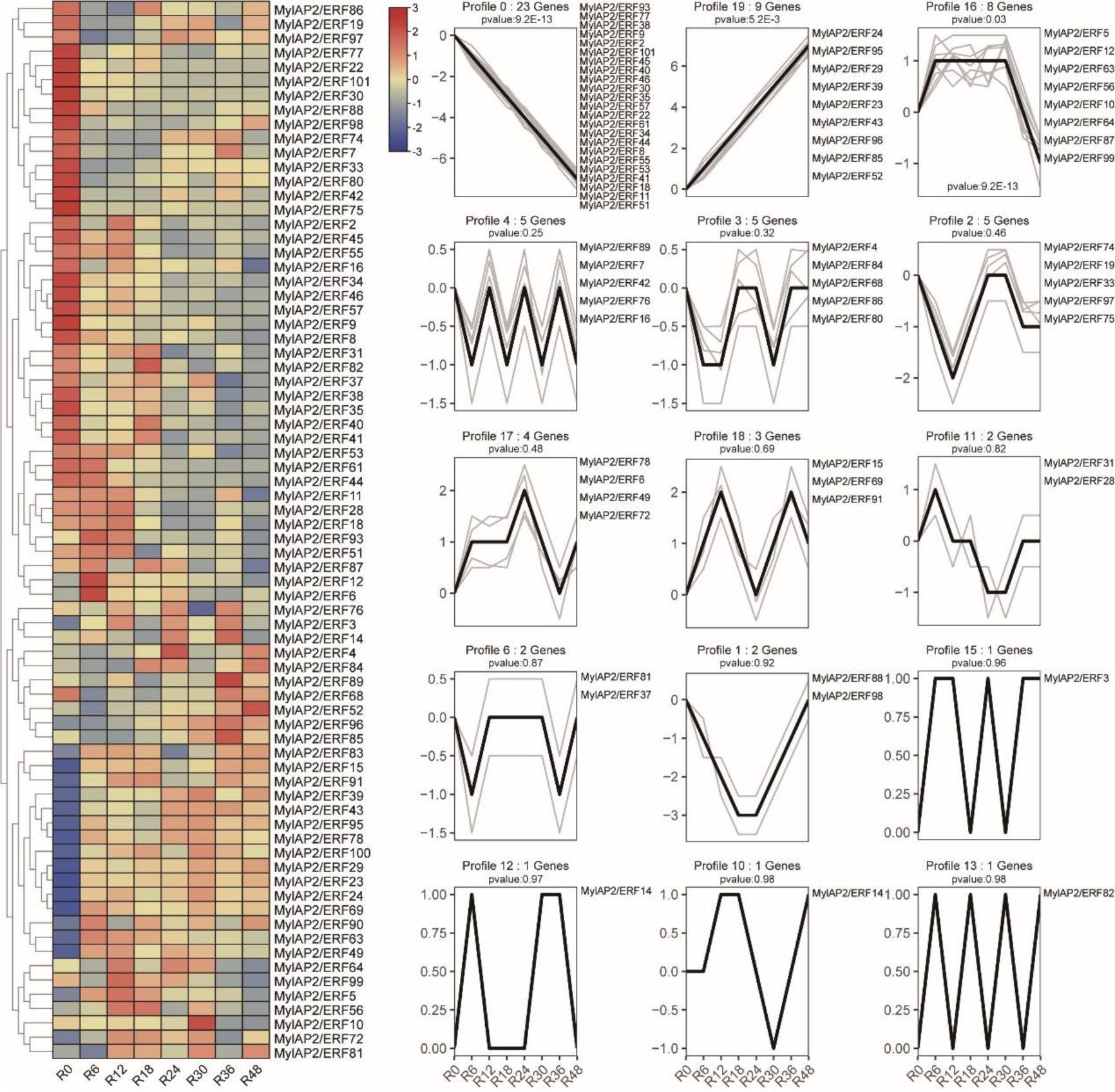
Expression patterns of MylAP2/ERF in water flooding stress.

## 3. Discussion

High quality reference genomes are of great significance for the basic biological research of a species [34,35]. With the continuous updates of sequencing technology, reference genomes of more and more species have been assembled one after another [36]. *M.laxiflora*, as a unique stress resistant and flood tolerant plant in the Yangtze River Basin of China, can be an ideal species for studying the genetic mechanism of plant response to flooding [6,7]. This study used a combination of second-generation sequencing data, HiFi sequencing, and Hi-C methods to obtain a reference genome with an N50 length of 29.5 Mb and a genome size of 1.29Gb. Therefore, the integration strategy used here has been proven to be very effective in assembling the genome of *M.laxiflora*. Meanwhile, a total of 23,666 protein coding genes were predicted within the reference genome using multiple strategies. These results provide basic information for subsequent research on *M.laxiflora*.

The AP2/ERF gene family is one of the important transcription factor families involved in plant growth, development, and response to various abiotic stresses [37]. Previous studies have shown significant differences in the number of AP2/ERF gene family members among different plants [38,39,40,41]. For example, 147 AtAP2/ERF members were found in Arabidopsis, 214 ZmAP2/ERF members were found in maize, and 543 AP2/ERF members were found in *Tritipyrum*. In this study, a total of 101 MylAP2/ERF members were found in the genome of *M.laxiflora*, which is relatively fewer compared to the aforementioned species. In addition, there are certain differences in the number of AP2/ERF subfamilies among different species. Most plants include four subgroups: AP2, RAV, ERF, and DREB. The ERF and DREB subfamily have a larger number of members, while the Soloist subgroup has a smaller number of members in plants. Among the 214 AP2/ERF members in maize, 166 (77.5%) members belong to the ERF and DREB subgroups, and no members of the Slolosist subgroup were found in *Tritipyrum*, rice, and maize [39,40,41]. In this study, only one Soloist member (MylAP2/ERF84) was found, while 80 members (79.2%) of the ERF and DREB subfamily were found. These results indicate that the number of AP2/ERF family members is mainly determined by the number of ERF and DREB subfamily members.

Whole genome replication is one of the main driving forces for the expansion of plant gene families, and is mainly achieved through two forms: tandem duplication and fragment duplication [42,43]. Previous studies have shown that the AP2/ERF gene family exhibits multiple whole genome replication events in different species. A total of 79 fragment duplication events were found in the AP2/ERF gene family of maize, and 32 tandem duplication events and 72 fragment duplication events were found in the AP2/ERF gene family study of *Tritipyrum* [40,41]. In *M.laxiflora*, a total of 10 tandem repeat events involving 21 MylAP2/ERF members and 32 fragment repeat events involving 47 MylAP2/ERF members were found. These implications suggest that compared to tandem repeat events, fragment repeat events may play a more important role in the expansion of the AP2/ERF gene family in plants.

Flooding stress is one of the abiotic stresses that plants suffer from, as they are exposed to cyclic or long-term hypoxic environments that limit their energy requirements for aerobic respiration and maintenance of life activities [45]. Multiple studies have shown that AP2/ERF members (*RAP2.2*, *HRE1*, *HRE2*) play important roles in the process of plant flooding stress [29,30]. Overexpression of Arabidopsis *RAP2.2* gene can significantly increase the expression abundance of hypoxia responsive genes (*PDC1* and *ADH1*), thereby significantly improving the survival rate of plants in hypoxic environments [29]. However, in the *rap2.2* mutant plants, the expression of *ADH1* and *PDC1* was not significantly reduced, suggesting that other ERF members or transcription factors may synergistically participate in plant hypoxia response [29]. Similarly, overexpression of *HRE1* and *HRE2* can quickly induce the expression of *ADH1* and *PDC1*, thereby improving plant hypoxia tolerance [30]. In this study, transcriptome data analysis at different time points of flooding stress showed that the expression abundance of 74 MylAP2/ERF members changed significantly. Among the 74 MylAP2/ERF members, the expression abundance of 23 MylAP2/ERF members continued to decline, while the expression abundance of 9 MylAP2/ERF members continued to increase Meanwhile, the results of phylogenetic analysis showed that MylAP2/ERF49, MylAP2/ERF78, MylAP2/ERF91 in *M.laxiflora*, and RAP2.2, HRE1, and HRE2 in Arabidopsis were in the same branch, and the expression abundance of MylAP2/ERF49 and MylAP2/ERF78 showed a trend of first increasing and then decreasing with the increase of flooding stress time, while the expression abundance of MylAP2/ERF91 showed a pattern of first increasing, then decreasing, and finally increasing again with the increase of flooding stress time. In summary, members of the AP2/ERF gene family play an important role in plant response to waterlogging stress and hypoxia stress.

*M.laxiflora* is one of the important plant species in the Yangtze River Basin in China, but its growth and development are often subjected to flooding stress, which poses a serious threat to its growth and development [3]. In recent years, with the development of sequencing technology and biotechnology, it is possible to further improve the waterlogging tolerance of *M.laxiflora* [2]. In this study, we assembled a high-quality reference genome for the first time, with a genome size of 1.29 Gb. At the same time, 101 AP2/ERF members were identified from the reference genome, and 74 MylAP2/ERF members were found to respond to flooding stress. In the future research, these AP2/ERF members can be precisely modified by biotechnology, so as to improve the waterlogging tolerance of *M.laxiflora*. Therefore, this study provides valuable information for further study on the biological function and molecular regulation mechanism of AP2/ERF gene in *M.laxiflora* and other plants.

## 4. Materials and methods

### 4.1 Plant materials

LS7, which was collected form Yangtze River Basin in China, was used to genome assembly in this study. To avoid sample contamination, asexual propagation of LS7 was carried out using cutting methods. Roots, leaves, stem, flowers and fruits were collected from 10 independent asexual propagation lines of LS7. Each sample was set with three replicates and quickly stored in liquid nitrogen for subsequent DNA and RNA sequencing.

### 4.2 Sequencing and Genome assembly

All sequencing libraries are constructed according to the quality requirements of sequencing equipment. The HiFi and Hi-C library was prepared according to the standard procedure [27]. RNA is isolated from different tissues (root, leaves, stem, flowers and fruits). The RNA-seq library was prepared using Illumina platform (Illumina, Santiago, CA, USA). A total of 16.99 Gb, 11.16 Gb, 10.24 Gb, 10.75 Gb and 11.39 Gb of RNA-seq raw data were obtained for root, leaves, stem, flowers and fruits, respectively. The raw reads were then filtered using SOAPfilter software.

K-mer analysis was employed to estimate the genome size. K-mers (K = 17) were counted by Jellyfish [29] using 93.0 Gb high-quality short reads.

After removing the HiFi reads of less than 1 Kb in length and adapters, 49.2 Gb clean reads, representing ∼ 44.7 × sequencing coverage of the *M.laxiflora* genome, were used for contig assembly with Falcon v0.3.0 [30]. Then, BLASR [31] was employed to map the reads to contigs. SMRT was used to correct some sequencing errors. BWA mem was used to map the paired-end reads without Illumina PCR to the corrected contig [32], and further use high-quality reads for assembly [33].

About 155.51 Gb of raw data was sequenced from a Hi-C library and ∼77.43 Gb of valid reads were obtained after quality control. The clean Hi-C reads were mapped to the genome assembly through BWA align. HiC-Pro was used for repetitive reading removal, classification and quality assessment [34]. Raw counts of Hi-C links were aggregated and separately by using Juicer [35] and 3D-DNA [36]. The Juicebox [37] software was used to adjust the placement and orientation errors.

### 4.3 Gene annotation

Gene prediction was performed by homology-based prediction, de novo prediction and transcriptome-based prediction. For homology-based prediction, protein sequences from eight species were mapped onto the *M.laxiflora* genome by an E-value cutoff of 10^−5^, and then Genewise was used for gene structure annotation. AUGUSTUS [44] (Version 2.03) and FGENESH [45] (Version 1.3) were used for de novo prediction. RNA-seq data from five tissues were processed by HISAT2 [46] and StringTie [47] for transcriptome-based prediction. EvidenceModeler [48] software was used to combined the above results and get the final non-redundant reference gene set. InterproScan [49], Gene Ontology (GO) [50], Kyoto Encyclopedia of Genes and Genomes (KEGG), SwissProt [51], TrEMBL and Non-redundant protein NCBI databases were used to annotate the functions of the predicted genes by BLAST searches (E-value cutoff 1 × 10^−5^). tRNAscan-SE [52] was used to identify tRNA. Rfam database and Infernal [53] were used to identify noncoding RNAs (including rRNA, miRNA and snRNA) based on homologous alignment. Both of De novo prediction and homology-based alignment were used to identify transposable elements. The De novo repeat database was built by a combination of the results of LTR_Finder [38], PILER [39], and RepeatScout [40]. This De novo repeat database together with Repbase [41] were used to identify repeats by RepeatMasker [42] and to identify repeat related proteins by RepeatProteinMask (http://www.repeatmasker.org/). RepeatMasker and TRF [43] were used to annotate the tandem repeats. The results above were combined according to their physical positions and transposable elements were further classified by blast against Repbase.

### 4.4 Identification and chromosomal distribution of AP2/ERF gene family in *M.laxiflora*

The AP2/ERF conserved domain PF00847 which obtained from the Pfam database (https://pfam.xfam.org/) was used to inquire about proteins in *M.laxiflora* reference genome [46]. Hidden Markov model (HMM) analysis with the parameter E-value < 1e-10 and bits > 85 were conducted to identify the candidate MylAP2/ERF members in the reference genome of *M.laxiflora* in this study. Finally, use the TBtools was used to visualize it on the chromosome position and name it based on its position on the chromosome. The cDNA sequence length, protein sequence length, molecular weight, and theoretical isoelectric points of identified MylAP2/ERF members were obtained from the GFF file of the *M.laxiflora* reference genome assembled in this study by using TBtools [47]. The subcellular localization of these MylAP2/ERF proteins were predicted by using the online tool WoLF PSORT (https://wolfpsort.hgc.jp/).

### 4.5 Phylogenetic analysis and motif analysis of MylAP2/ERFs

The MEGA11 software was used to conduct the phylogenetic analysis on all members of the AP2/ERF gene family in *M.laxiflora* and Arabidopsis. Before constructing the phylogenetic tree, use MEGA11 software to align the AP2/ERF protein sequences from *M.laxiflora* and Arabidopsis, with default parameters set. Using the NJ method to construct phylogenetic trees for MylAP2/ERF and AtAP2/ERF, the specific parameters are the Poisson model and the Bootstrap duplicate value set to 1000, while the rest are default parameters. The conserved motif analysis among the MylAP2/ERF members was conducted by using the MEME online program (https://meme-suite.org/meme/index.html) and set the parameters as default, with a maximum motif number of 10.

### 4.6 Gene duplication event and *cis*-acting elements analysis

Identification and analysis of gene duplication events in MylAP2/ERF gene family were performed by using MCScanX software, and the results were visualized using TBtools [47, 48]. Extract the upstream 2000 bp sequence of the MylAP2/ERF gene start codon (ATG) as the promoter region by using TBtools, and the online website PlantCARE (https://bioinformatics.psb.ugent.be/webtools/ Plant care/HTML/) was used to predict the *cis*-acting elements in the promoter regions of these MylAP2/ERF members.

### 4.7 Expression analysis of MylAP2/ERF genes

The expression analysis of these MylAP2/ERF genes include the different tissue expression patterns and under flooding stress expression patterns. The transcriptome data for different tissues (leaves, flowers, stems, and fruits) of *M.laxiflora* were obtained in this study. And, the transcriptome data for *M.laxiflora* roots under flooding stress at different time points was download from the public database NCBI (PRJNA840865). Gene expression levels were calculated based on the fragments per kilobase of exon per million mapped fragments (FPKM) value. And, differentially expressed genes were retrieved using DESeq with the following parameters: padj < 0.05 and |log2FC| ≥ 1 [49]. Finally, the heat map of all relevant data was drawn using the TBtools [47].

## 5. Conclusion

Here, we report for the first time the high-quality reference genome of *M. laxiflora* by using HiFi sequencing and Hi-C. The total assembly size of the *M. laxiflora* genome was 1.29 Gb with a scaffold N50 size of 29.5 Mb. A total of 23,666 genes encoding proteins and 5,457 ncRNA were predicted in this reference genome. In the *M. laxiflora* genome, a total of 101 MylAP2/ERF genes were identified and divided into 5 subgroups: 40 belong to the ERF subgroup, 40 belong to the DREB subgroup, 17 belong to the AP2 subgroup, 3 belong to the RAV subgroup and 1 belongs to the Soloist subgroup. And, a total of 10 tandem repeat events and 32 fragment duplication events were found among the 101 MylAP2/ERF genes. A large number of MeJA responsive elements, ABA responsive elements, and hypoxia responsive elements were found in the promoter regions of these MylAP2/ERF genes. Based on transcriptome data analysis, a total of 74 MylAP2/ERF genes can respond to flooding stress. Meanwhile, it was found that three genes (MylAP2/ERF49/78/91) which belong to the same branch with RAP2.2 gene exhibited different expression trends under flooding stress. Our results provide valuable information on the molecular regulatory mechanism of flooding stress in *M. laxiflora*.

## Data availability

The final chromosome-level genome and annotation files of *Myricaria laxiflora* is available in figshare, doi: 10.6084/m9.figshare.25375366.v1.

## Additional files

See Supplementary data

## Authors’ contributions

Bicheng Dun, Huiyuan Chen, Xiaobo Ma collected the plant materials; Guiyun Huang, Di Wu, Jihong Liu designed the project; Weibo Xiang, Linbao Li, Zhen Yang, Zhiqiang Xiao and Haibo Zhang worked on sequencing and data analysing; Weibo Xiang and Linbao Li wrote the manuscript; All the authors revised and approved the final version of the manuscript.

